# Exploration of social spreading reveals behavior is prevalent among *Pedobacter* and *P. fluorescens* isolates, and shows variations in induction of phenotype

**DOI:** 10.1101/758656

**Authors:** Lucy M. McCully, Jasmine Graslie, Adam S. Bitzer, Auður M. Sigurbjörnsdóttir, Oddur Vilhelmsson, Mark W. Silby

**Author notes:** For correspondence: Mark Silby, Department of Biology, University of Massachusetts Dartmouth, 285 Old Westport Road, North Dartmouth, MA 02747. These authors contributed equally to this work (Lucy M. McCully, Jasmine Graslie). Submit to: AEM, FEMS Micro Letters, Microbiology SGM.

## Abstract

Within soil, bacteria are naturally found in multi-species communities, where interactions can lead to emergent community properties. It is critical that we study bacteria in a social context to investigate community-level functions. We previously showed that when co-cultured, *Pseudomonas fluorescens* Pf0-1 and *Pedobacter* sp. V48 engage in interspecies social spreading on a hard agar surface, a behavior which required close contact and was dependent on the nutritional environment. In this study, we investigate whether the ability to participate in social spreading is widespread among *P. fluorescens* and *Pedobacter* isolates, and whether the requirements for interaction vary. We find that this phenotype is not restricted to the interaction between *P. fluorescens* Pf0-1 and *Pedobacter* sp. V48, but is a more prevalent behavior found in one clade in the *P. fluorescens* group and two clades in the *Pedobacter* genus. We also discovered that the interaction with certain *Pedobacter* isolates occurred without close contact, indicating induction of spreading by a putative diffusible signal. As is the case for ISS by Pf0-1+V48, motility of all interacting pairs is influenced by the environment, with no spreading behaviors observed under high nutrient conditions. While Pf0-1+V48 require low nutrient but high NaCl conditions, in the broader range of interacting pairs this requirement for low nutrient and high salt was variable. The prevalence of motility phenotypes observed in this study and found within the literature indicates that community-induced locomotion in general, and social spreading in particular, is likely important within the environment. It is crucial that we continue to study microbial interactions and their emergent properties to gain a fuller understanding of the functions of microbial communities.

## Introduction

Within every soil environment exist rich communities with members from not only a variety of microbial species, but inhabitants representing every domain of life. Microbes are ubiquitously present within soil environments, with microbial communities playing important roles within this ecosystem (1, 2). The soil microbiome is required for biogeochemical cycles, processes that regulate availability of phosphorus, nitrogen, carbon, and other nutrients, and can also affect plant health through pathogen defense, nutrient uptake, and growth (1, 2). These influences on both abiotic and biotic factors can have ecosystem-level effects, mediating nutrient cycling, bioremediation, and even altering the composition of the aboveground plant communities (3–5).

There exist important functions of microbial communities that cannot be attributed to any one taxa, and are, instead, a property of the collective. These emergent properties result from social interactions between different microbial species in a community context, and, importantly, cannot be predicted based on the study of the individual microbes in isolation (6, 7). These community-intrinsic properties emerge from the interactions within a community, which can be as small as a pair of microbial species (8). Emergent traits can affect large-scale ecosystem processes in ways we do not yet understand, but which may be revealed by studying groups of microbes in co-culture. Experimental studies of microbial interactions, particularly mechanistic studies, are needed to improve our understanding of the emergent properties of communities, and microbial ecology (7, 8).

Altered motility has been documented in several studies as an emergent phenotype from interspecies interactions, often as a prevalent phenotype between two groups of microbes. To escape competition, *Bacillus subtilis* was found to engage in sliding motility away from various species of antibiotic-producing *Streptomyces* antagonists (9, 10). *Streptomyces venezuelae* itself has been shown to engage in ‘exploratory’ motility across an agar surface in response to the presence of yeast, and a number of other fungal species (11). Among 200 other *Streptomyces* isolates tested, 19 exhibited a similar exploratory behavior, but these isolates did not form a monophyletic group. Limoli et al. showed a behavior where *Pseudomonas aeruginosa* responds to the presence of *Staphylococcus aureus*, by using its pili to engage in ‘exploratory’ motility towards a *S. aureus* microcolony (12). Clinical isolates of both species were shown to induce (*S. aureus*) or respond with exploratory behavior (*P. aeruginosa*), indicating a more widespread phenomenon. *Xanthomonas perforans*, as well as various other *Xanthomonas* isolates, were found to induce surface motility in swarming *Paenibacillus vortex*, in order for *Xanthomonas* to hitchhike on top of the *P. vortex* swarming rafts (13). *X. perforans* was also found to hitchhike on other *Paenibacillus* isolates, as well as *Proteus mirabilis*. These studies highlight the possibly critical importance of finding a way to move within the natural environment. The ability to engage in these altered behaviors broadens capacity for motility within communities, beyond the mechanisms and triggers characterized in individual species.

We previously showed that the mixture of two soil bacteria, *Pseudomonas fluorescens* Pf0-1 and *Pedobacter* sp. V48, led to a spreading behavior across a hard agar surface (14). These two species were not motile individually, under these culture conditions, indicating that the consortium only moves when the two species are together. Through microscopy and culturing methods, we demonstrated that the two species co-migrate, and that close proximity is required for induction of movement, as neither preconditioning the agar with the other species, or addition of dead cells could induce motility in the partner species. We also found that the interaction is conditional; no motility behavior was observed under high nutrient conditions, and under relatively low nutrient conditions, the addition of salt was required to observe the social spreading phenotype. The lack of motility observed in each species individually, along with the requirement of the physical presence of both species for the social spreading phenotype places this behavior in contrast with previously-described socially-derived motile phenotypes, which require at least one of the species involved to be individually motile under the experimental conditions (13, 15, 16)

This previous research raised the question about whether interspecies social spreading could also be demonstrated in related isolates of both partner species, and which characteristics would be conserved (14). We hypothesized that closely-related species would exhibit similar motility phenotypes, and that uncovering these phenotypes would lead to both a better understanding of the motility mechanism, as well as its environmental relevance. We gathered additional *P. fluorescens* and *Pedobacter* isolates to test in the interaction assay with the partner species. We demonstrate that a monophyletic group of each species exhibited the same behavior as the canonical Pf0-1+V48 interaction, and that a second *Pedobacter* clade induces motility in Pf0-1 through a diffusible signal. We also show that no interactions occur between any species combinations in high nutrient conditions, and see variability in the degree of surface spreading in low nutrient conditions with and without raised salt content.

## Results

### Motility phenotypes similar to interspecies social spreading are found among closely-related isolates

To explore the breadth of related isolates that are capable of participating in interspecies social spreading, we tested 17 additional *Pedobacter* species for their interaction with *P. fluorescens* Pf0-1, and 18 additional isolates of the *P. fluorescens* group for their interaction with *Pedobacter* sp. V48. To correlate the phenotypic similarities of the isolates capable of social spreading with degrees of relatedness, we constructed phylogenetic trees (Figures 1, 2). The more distantly related *Pseudopedobacter saltans* and *Pseudomonas aeruginosa* PAO1, which serve as outgroups in the phylogenetic trees, were also tested in the social assay.

**Figure 1.**
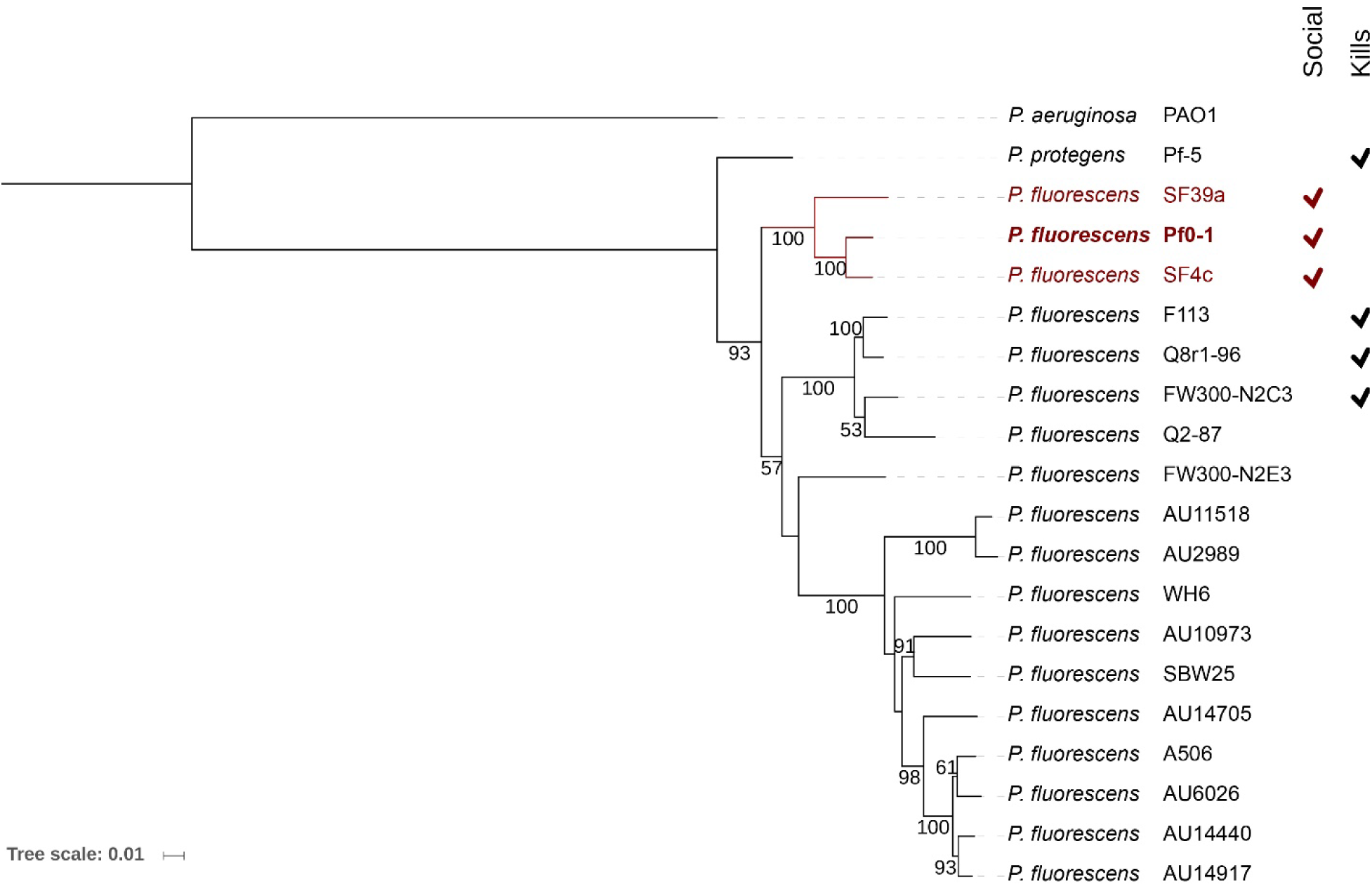
Phylogenetic relationship of the *P. fluorescens* group, based on housekeeping gene sequences (*gapA, gltA, gyrB, and rpoB)*. The evolutionary history was inferred by using the Maximum Likelihood method based on the General Time Reversible model (17) using MEGA7 (18). Numbers at branch nodes indicate confidence levels (bootstrap probability inferred from 1000 replicates) greater than 50%. All species were tested by co-culture with *Pedobacter* sp. V48, with those exhibiting interspecies social spreading behavior in dark red, and a column for those that kill *Pedobacter* sp. V48. *P. fluorescens* Pf0-1 is indicated in bold. *P. aeruginosa* PAO1 was used as an outgroup. Scale bar, 0.01 substitutions per nucleotide position.

**Figure 2.**
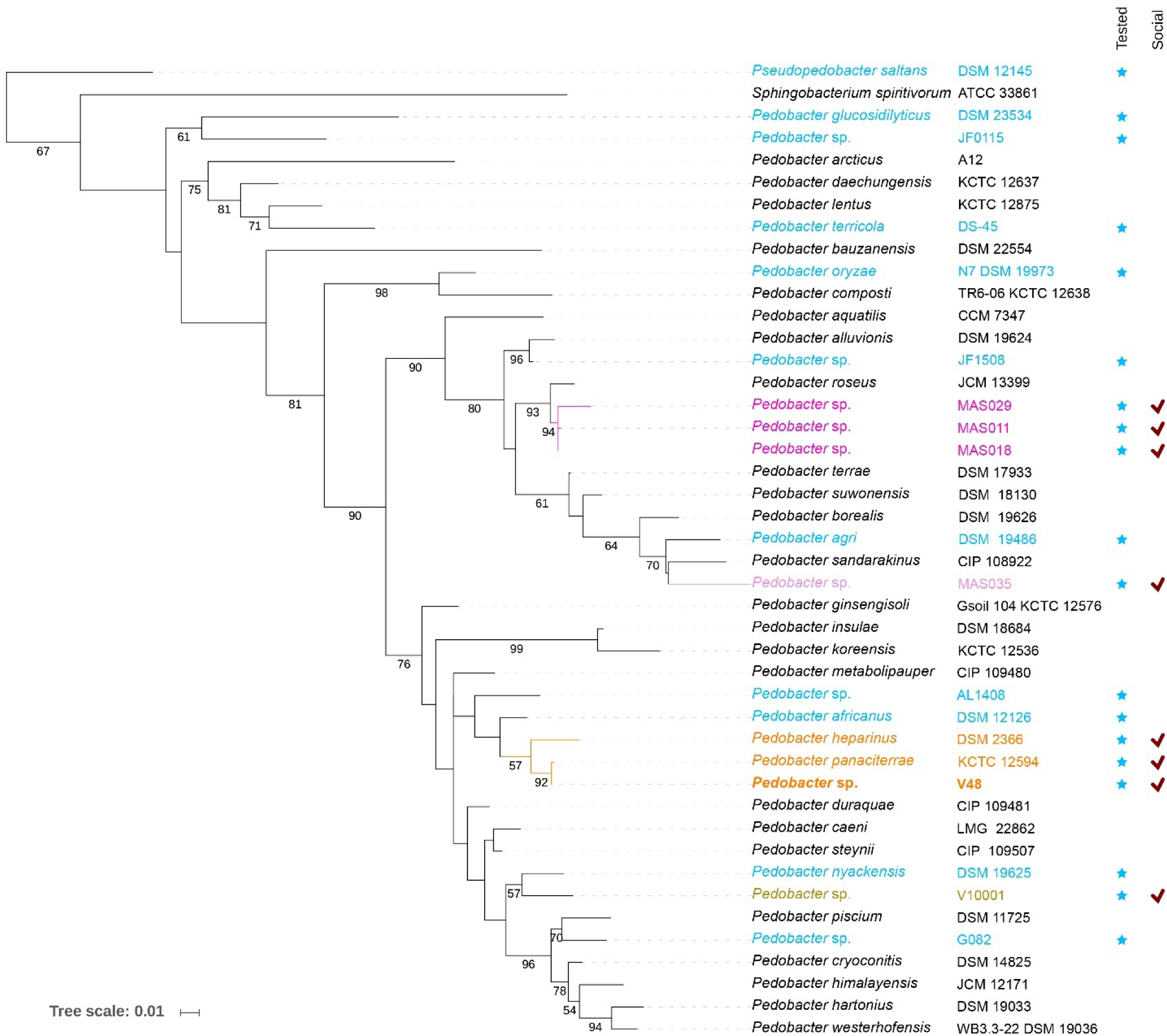
Phylogenetic relationship of *Pedobacter* species, based on 16S rRNA gene sequences. The evolutionary history was inferred by using the Maximum Likelihood method based on the General Time Reversible model (17) using MEGA7 (18). Numbers at branch nodes indicate confidence levels (bootstrap probability inferred from 1000 replicates) greater than 50%. In blue are shown species tested by co-culture with *P. fluorescens* Pf0-1, with those exhibiting different kinds of social spreading in pink (MAS clade), light pink (MAS035), orange (V48 clade), and olive (V10001). *Pedobacter* sp. V48 is indicated in bold. *Pseudopedobacter saltans* DSM 12145 was used as an outgroup. Scale bar, 0.01 substitutions per nucleotide position.

We observed the ability to participate in social spreading behavior among a subset of isolates from both *Pedobacter* and *Pseudomonas* (Figure 3). *P. fluorescens* SF4c and SF39a, the two isolates clustered closest to Pf0-1 (Figure 1), were also those that exhibited social spreading behavior with *Pedobacter* sp. V48 (Figure 3B, C). The spreading patterns were similar, but not identical, to those of social spreading by *P. fluorescens* Pf0-1 and *Pedobacter* sp. V48 (Figure 3A). SF4c social motility has a distinct crinkled appearance (Figure 3B), while SF39a expands asymmetrically, with a smooth colony and lobate edges (Figure 3C). As is the case in Pf0-1 interaction with V48, SF4c and SF39a each co-migrate with V48, as evidenced by isolation of both partners from both the middle and the edge of the spreading colony.

**Figure 3.**
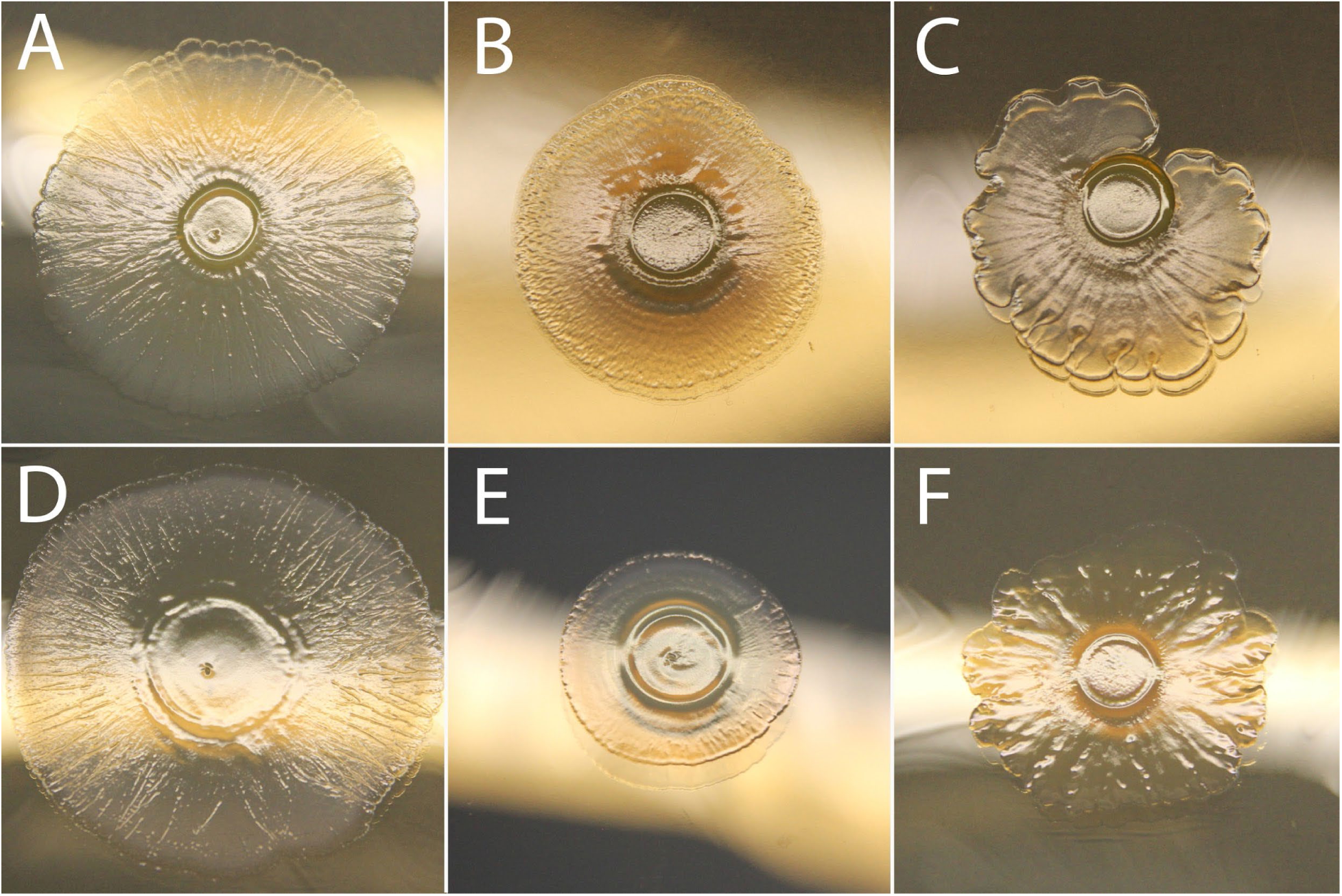
Patterns resulting from spreading colonies of mixed co-cultures of different *P. fluorescen*s and *Pedobacter* isolates. A) *Pedobacter* sp. V48 and *P. fluorescens* Pf0-1, B) *Pedobacter* sp. V48 and *P. fluorescens* SF4c, C) *Pedobacter* sp. V48 and *P. fluorescens* SF39a, D) *Pedobacter panaciterrae* and *P. fluorescens* Pf0-1, E) *Pedobacter heparinus* and *P. fluorescens* Pf0-1, F) *Pedobacter* MAS011 and *P. fluorescens* Pf0-1 (representative of MAS018 and MAS029). Colonies were incubated 144 h.

The *Pedobacter* isolates capable of social spreading are not randomly distributed, but rather group into two distinct clades (Figure 2), each displaying distinct social spreading patterns. The clade we refer to as the V48 clade, clustering most closely with *Pedobacter panaciterrae* and *Pedobacter heparinus*, had similar patterns of social spreading to *Pedobacter* sp. V48 when paired with Pf0-1 (Figure 3A, D, E), though the degree of expansion differed over the same period of time. Each species was found in the middle and at the edge of the spreading colony, indicating co-migration with Pf0-1. The other social-spreading clade, consisting of *Pedobacter* MAS011, MAS018, and MAS029 and thus termed the MAS clade, exhibited an interspecies social spreading pattern that differed from V48, with an irregular, undulate edge, and a more mucoid mixed colony (Figure 3F). It also differed in that all the MAS mono-cultures exhibit a sporadic and modest amount of irregular colony expansion with feather-like patterns. The MAS clade was found to cluster most closely with *Pedobacter roseus* (Figure 2).

### A distinct *Pedobacter* clade interacts with Pf0-1 without close association: MAS011, MAS018, and MAS029

We previously demonstrated that close association was required for the social spreading of Pf0-1 and V48 (14). To ask whether this association was a general requirement for social spreading between Pf0-1 and *Pedobacter* partners, we plated Pf0-1 colonies adjacent to other interacting *Pedobacter* isolates (identified above). Consistent with *Pedobacter* sp. V48 (14), *P. panaciterrae* and *P. heparinus* required physical contact to engage in the social phenotype. In contrast, five other *Pedobacter* isolates were able to trigger surface spreading in Pf0-1 when growing without contact between the species (Figure 4). Spreading of Pf0-1 begins asymmetrically, at the colony face closest to the *Pedobacter* isolate. Once spreading becomes more extensive, the direction of spreading often homogenizes so the Pf0-1 colony appears to spread in all directions (Figure 4). The *Pedobacter* isolates MAS011, MAS018, MAS029, and MAS035 were able to induce motility from a distance of approximately 14 mm, while V10001 induced at up to 10 mm, although this strain was not tested at a narrower range (between 10 mm and 15 mm). In addition to Pf0-1 being induced to move, a reciprocal effect on MAS clade *Pedobacter* isolates (MAS011, MAS018, and MAS029) was frequently observed. The members of the MAS clade (Figure 2) frequently exhibited a feather-like spreading phenotype in the presence of Pf0-1 (Figure 4A). In contrast with the V48 clade, *Pedobacter* isolates from the MAS clade *can* be isolated out to the edge of the colony, but are not consistently found there, suggesting co-isolation is coincidental, and they migrate independently.

**Figure 4.**
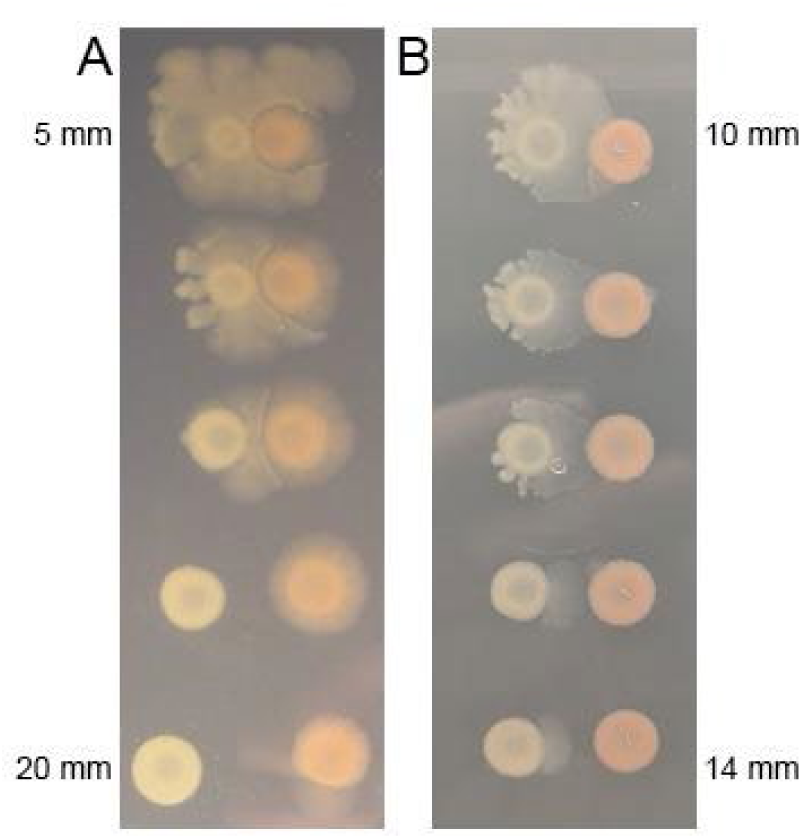
*P. fluorescens* Pf0-1 surface spreading in response to *Pedobacter* strain MAS011 putative diffusible signals. A) *P. fluorescen*s Pf0-1 (left) and *Pedobacter* MAS011 (right) inoculated at distances of 5, 7.5, 10, 15, and 20 mm from top to bottom, B) *P. fluorescen*s Pf0-1 (left) and *Pedobacter* MAS011 (right) inoculated at distances of 10 mm to 14 mm, with 1 mm intervals. Plates inoculated for 7 days.

### Social spreading in the interaction of Pf0-1 and the *Pedobacter* MAS clade is mediated by putative diffusible, but not volatile, signals

Because *Pedobacter* isolates MAS011, MAS018, MAS029, and MAS035 induce motility of Pf0-1 at a distance, we asked whether this interaction was mediated by a diffusible or volatile signal. Agar was pre-conditioned by first growing MAS011, MAS018, MAS029, or MAS035 (individually) on a porous cellophane membrane. Pf0-1 plated after removal of cellophane and *Pedobacter* cells was able to spread alone across the agar surface whereas Pf0-1 plated on plates not pre-treated with *Pedobacter* isolates did not spread. There was a significant difference in diameter when comparing treated and control colonies (p < 0.05), suggesting secretion of a signal for motility by the *Pedobacter* MAS isolates.

We also asked whether this motility in Pf0-1 could be induced by volatile signals, as *P. fluorescens* Pf0-1 and *Pedobacter* sp. V48 have previously been shown to interact through both diffusible and volatile signals (19–21). Pf0-1 and MAS011, MAS018, MAS029, and MAS035 isolates were plated alone in the center of TSB-NK plates. The plates were sandwiched together in pairs, each pair containing Pf0-1 and a MAS isolate. Pf0-1 did not display a motility phenotype over the course of 7 days with any of the observed MAS isolates (change in diameter not significant).

### Differences in social spreading phenotypes within other clades: MAS035 and V10001

In addition to the two clades with distinct motility triggers (diffusible signals vs close contact), we observed two phylogenetically isolated *Pedobacter* isolates that induce surface spreading in *P. fluorescens* Pf0-1 (Figure 5). *Pedobacter* MAS035 exhibited a near-identical social spreading phenotype to the other three MAS isolates (Figure 3F). However, in contrast to the members of that clade, MAS035 was never culturable from the edge of the moving colony. Closer examination revealed that the pink pigment of MAS035 was only visible in the center (Figure 5D), suggesting the isolate remains localized at the point of inoculation. As with the other isolates of the MAS clade, Pf0-1 migrates when plated at a distance from MAS035 (Figure 5C). However, *Pedobacter* MAS035 never develops the feather-like spreading phenotype of the MAS clade in the presence of Pf0-1 (Figure 5B, 4A). *Pedobacter* V10001, like MAS035, induces migration in Pf0-1 before the colonies come into contact (Figure 5G). In a striking contrast with other *Pedobacter-*Pf0-1 interactions, when Pf0-1 is co-cultured (mixed) with V10001, the colony remains immotile (Figure 5H). An explanation for this phenomenon remains elusive.

**Figure 5.**
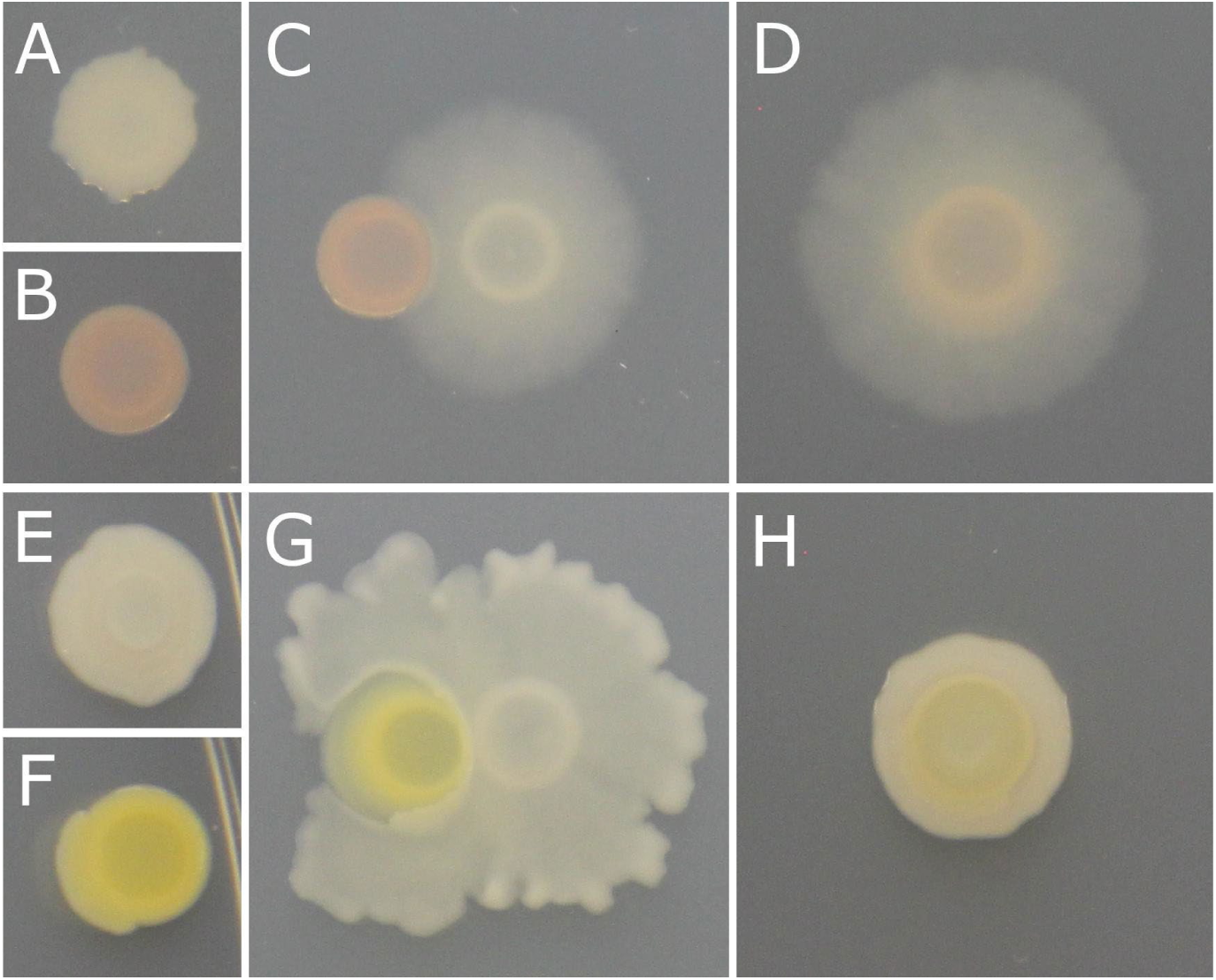
Motility phenotypes within phylogenetically separate *Pedobacter* isolates. A) *P. fluorescens* Pf0-1 mono-culture, B) *Pedobacter* MAS035 mono-culture, C) *Pedobacter* sp. MAS035 (left) and *P. fluorescens* Pf0-1 (right) plated adjacently, D) *Pedobacter* sp. MAS035 and *P. fluorescens* Pf0-1 co-culture, E) *P. fluorescens* Pf0-1 mono-culture, F) *Pedobacter* V10001 mono-culture, G) *Pedobacter* sp. V10001 (left) and *P. fluorescens* Pf0-1 (right) plated adjacently, H) *Pedobacter* sp. V10001 and *P. fluorescens* Pf0-1 co-culture. Colonies in panels A, B, C, and D were incubated for 72 hours. Colonies in panels E, F, G, and H were incubated for 144 hours.

### Interactions between various *Pedobacter* and *Pseudomonas* isolates do not replicate the same behaviors as the interactions with V48 and Pf0-1

We next asked if interactions would occur between the motile *P. fluorescens* and the motile *Pedobacter* isolates we had tested previously (i.e. isolates from each genus/group other than V48 and Pf0-1). We tested *P. fluorescens* SF4c and SF39a in the mixed culture and side-by-side interaction assays with isolates from the V48 clade, *P. panaciterrae* and *P. heparinus*, isolates from the MAS clade, *Pedobacter* MAS011, MAS018, and MAS029, as well as the phylogenetically independent isolates, *Pedobacter* MAS035 and V10001.

Neither *P. fluorescens* strains recapitulate the Pf0-1 phenotypes when tested with the two other species within the V48 clade, but do show interesting interaction-associated spreading (Table 1). Neither of the strains were found to interact with *P. heparinus*. When plated adjacent to the *P. panaciterrae* colony, the *P. fluorescens* strains show an individual spreading phenotype. Interestingly, co-migrating sectors consistently emerge from the mixed and adjacent colonies of *P. fluorescens* SF4c and *P. panaciterrae* (Figure 6), but do not develop until after plates have been incubating for eight days when adjacent, or after 7-10 days when mixed. In one instance, a co-migrating sector emerged from a mixed colony of *P. fluorescens* SF39a and *P. panaciterrae*.

**Table 1.** Social spreading of different combinations of *Pedobacter* and *P. fluorescens* isolates.

|  | *P. fluorescens* Pf0-1 |  |  | *P. fluorescens* SF4c |  |  | *P. fluorescens* SF39a |  |  |
| --- | --- | --- | --- | --- | --- | --- | --- | --- | --- |
|  | Mixed | Adjacent colonies | Migrate together<sup>1</sup> | Mixed | Adjacent colonies | Migrate together<sup>1</sup> | Mixed | Adjacent colonies | Migrate together<sup>1</sup> |
| *Pedobacter* sp. V48 | yes | together | yes | yes | together | yes | yes | together | yes |
| *Pedobacter panaciterrae* | yes | together | yes | no* | SF4c/yes** | Yes*** | no<sup>#</sup> | SF39a | no<sup>#</sup> |
| *Pedobacter heparinus* | yes | together | yes | no | no | no | no | no | no |
| *Pedobacter* MAS011 | yes | both, without contact | sometimes<sup>^</sup> | yes | both, without contact | no | yes | both, without contact | no |
| *Pedobacter* MAS018 | yes | both, without contact | sometimes<sup>^</sup> | yes | both, without contact | no | yes | both, without contact | no |
| *Pedobacter* MAS029 | yes | both, without contact | sometimes<sup>^</sup> | yes | both, without contact | no | yes | both, without contact | no |
| *Pedobacter* MAS035 | yes | Pf0-1 | no | yes | SF4c | no | yes | SF39a | no |
| *Pedobacter* V10001 | no | Pf0-1 | no | no | SF4c | no | no | SF39a | no |

<sup>1</sup>Indicates whether there is co-migration in mixed colonies. "No" indicates isolation of only *P. fluorescens* at the edge of the moving colony
<sup>^</sup> *Pedobacter* not consistently isolated at colony edge
\* except for sectors that emerge after plates have been incubating for 5-10 days
\*\* SF4c colony is individually motile at first, then a co-motile front emerges after 8 days
\*\*\* *Pedobacter* can be isolated from edge of the motile sector, but not usually at the edge of non-motile areas of the colony
<sup>#</sup>With the exception of one sector that emerged after 96 hours of incubation

**Figure 6.**
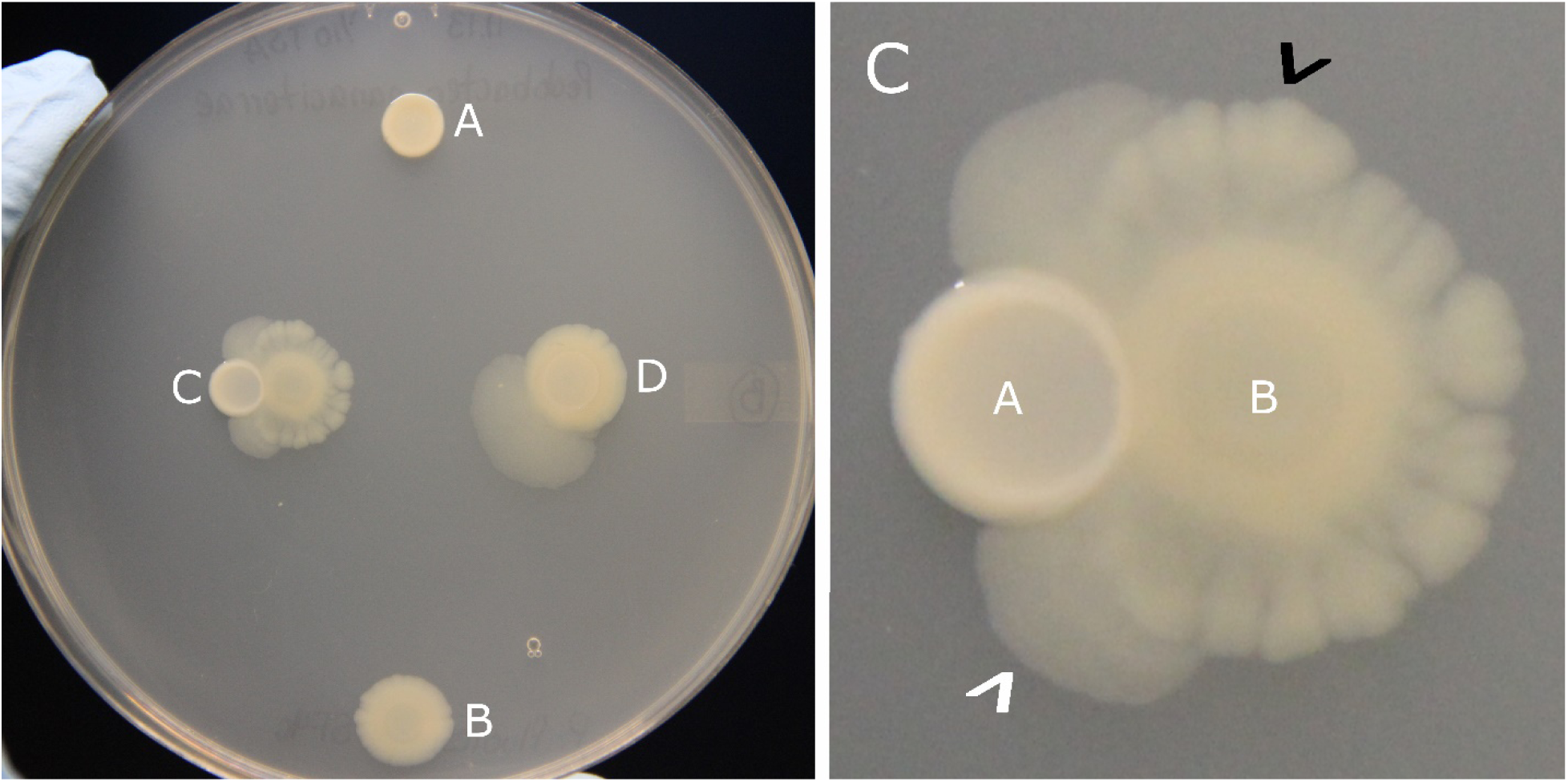
Plate showing *Pedobacter panaciterrae* and *P. fluorescens* SF4c incubated for 13 days. A) *P. panaciterrae* mono-culture, B) *P. fluorescens* SF4c mono-culture, C) Side-by-side colonies of *P. panaciterrae* (left) and *P. fluorescens* SF4c (right), D) Mixed colony of *P. panaciterrae* and *P. fluorescens* SF4c, showing the co-migrating sector that emerged from the ‘main’ colony C) Zoomed in inset showing the thicker individual spreading of *P. fluorescens* SF4c (black arrowhead) and the co-migrating motility that emerged on either side of the connected colonies after eight days (white arrowhead). Co-migrating motility developed after the plates had been incubating for eight days. Plate representative of three replicates.

Similar behavior to Pf0-1 was found when both *P. fluorescens* SF4c and SF39a were tested with the *Pedobacter* MAS isolates (Table 1). These two *P. fluorescens* strains spread when adjacent to or mixed with MAS011, MAS018, MAS029, and MAS035. However, unlike with Pf0-1, the three *Pedobacter* MAS isolates within the same clade never co-migrate with these *P. fluorescens* isolates when plated as a mixed culture. In side-by-side assays, the three MAS isolates also spread in the presence of the *P. fluorescens* isolates, though more modestly with SF39a. *P. fluorescens* SF4c and SF39a interact with *Pedobacter* V10001 in same way as Pf0-1 does (Table 1), as the colonies spread when adjacent to V10001, but not when cultures are mixed.

### Nutritional environment influences social spreading

We previously reported that the social spreading between Pf0-1 and V48 was influenced by the nutritional environment. Under high nutrient conditions, no social spreading occurred. Under reduced nutrient conditions, social spreading was slow, but could be accelerated in media with additional NaCl (5 g/L) (14). We tested Pf0-1 with *Pedobacter* isolates which we show above can partner in social spreading, and tested V48 with *P. fluorescens* strains we show above to be partners in social spreading (Table 1). In each interaction tested here, a high nutrient condition (TSB, 100%) precludes any social spreading phenotypes. The interactions of Pf0-1 with *P. heparinus* as well as SF4c with V48 both follow the same pattern as observed for Pf0-1 with V48 (14). However, social spreading among the remaining interacting pairs varied considerably (Table 2). Pf0-1 tested with MAS011, MAS018, MAS029 and MAS035 spreads regardless of the amount of NaCl, although this spreading is slower with additional NaCl in the medium. Similarly, V10001 and Pf0-1 spread regardless of NaCl addition when mixed or plated adjacently, in contrast with the behavior observed on TSB-NK (Table 1). Pf0-1 and *P. panaciterrae* spread in low nutrient conditions with and without added NaCl, as do SF39a and V48; the former pair is notably faster under conditions with high salt, while the latter pair is exceptionally rapid under conditions with no additional NaCl.

**Table 2.** Social spreading under different nutrient conditions.

|  |  | High<br>(TSB, 100%) | Low<br>(TSB-N) | Low<br>(10% TSB) |
| --- | --- | --- | --- | --- |
| *P. fluorescens* | *Pedobacter* |  | Added<br>NaCl | No added<br>NaCl |
| Pf0-1 | V48 | - | ++ | - |
| SF4c | " | - | + | - |
| SF39a | " | - | + | ++ |
| Pf0-1 | *P. panaciterrae* | - | ++ | + |
| " | *P. heparinus* | - | + | - |
| " | MAS011 | - | + | ++ |
| " | MAS018 | - | + | ++ |
| " | MAS029 | - | + | ++ |
| " | MAS035 | - | + | ++ |
| " | V10001 | - | + | + |

- non-motile
+ social spreading across a surface, but slow
++ social spreading across a surface

## Discussion

Interspecies social spreading was initially discovered as a surface spreading phenotype resulting from the interaction of *P. fluorescens* Pf0-1 and *Pedobacter* sp. V48 (14). The discovery of this phenotype raised the important question of the prevalence of similar motility behaviors among related species. In this study we show that interactions between various *Pedobacter* sp. and fluorescent pseudomonads can lead to the movement of a two-species consortium across a surface, although that ability is not conserved across the entire phylogenetic tree. The social behaviors we found appear similar, but there is some degree of variability in the phenotypes and modes of spreading (for example, whether or not motility is induced by diffusible signals), where closely-related strains appear to share more similar phenotypes. We observe that the extent of social spreading varies between strains, as well as between nutrient and salt content of the media, indicating an interplay between genetic contribution and environmental conditions.

The observed phenotypes display a common theme: the ability for the community to spread across a hard agar surface. The individual species don’t move in isolation under the conditions used, but as part of a group they have the ability to locomote. The result of the interaction between many of the isolates tested is motility, regardless of the variety in phenotypes, with differences such as motility patterns, how motility is induced, and possibly even the mechanism through which motility is achieved. There are a variety of examples of motility induced by culturing different species together; whether they appear cooperative, such as consortia gaining the ability to traverse environments each species could not cross individually (15, 22, 23), or competitive, such as *P. aeruginosa* ‘exploring’ towards *S. aureus* microcolonies to invade them, *B. subtilis* responding to competitors by sliding away from the antibiotics they produce, or *Streptomyces* exploratory motility to overtake environments containing a yeast competitor (9–12). The outcome for many bacterial interactions is movement, which underscores the importance of motility for many bacteria as a response to environmental challenges. The expanding number of examples of interaction-associated motility means the concept of motility in natural environments needs to be considered more broadly than just in terms of whether individual members can move in mono-culture in laboratory conditions. The role and prominence of motility in the function and ecology of microbial communities needs to explored with this in mind.

Our initial work characterized the social spreading phenotype resulting from the interaction of two soil bacteria, the distantly-related *P. fluorescens* Pf0-1 and *Pedobacter* sp. V48 (14). In this study, we have demonstrated that interactions between distantly-related strains leading to motility phenotypes are more prevalent than for just these two strains. We tested 18 *Pedobacter* species and 19 *P. fluorescens* strains for interspecies social spreading, and found that relatedness is an important factor, as the most closely related isolates have the same phenotypes, indicating there may be a genetic contribution from both partner bacteria dictating the observed phenotypes which is conserved among close relatives. Although the *Pedobacter* phylogenetic tree is derived from alignment of 16S sequences, its structure is generally consistent with the literature (24–32), giving us a higher degree of confidence in the relationships between the isolates. Interestingly, co-isolation of strains did not appear to be an important factor for their ability to engage in a social behavior, and even our canonical two species were not co-isolated, one having been isolated in Dutch dune soil, and the other from sandy-loam soil in a farm in MA, U.S.A (33, 34). The vast majority of the species we tested were isolated from soil, but *Pseudomonas* and *Pedobacter* can be isolated from a variety of different environments (35–40). Ultimately, close relationship to our initial two study strains is a better predictor of interaction than the environment from which the strains were isolated.

Our experiments plating different isolates at a distance, as well as those using semi-permeable membranes to precondition the agar, suggest the presence of a secreted diffusible signal from various *Pedobacter* species that we don’t observe in the canonical interaction between *P. fluorescens* Pf0-1 and *Pedobacter* sp. V48 (14). This indicates that direct contact is not required to induce motility in other interactions. The asymmetry we observe in the movement, where Pf0-1 colony begins spreading in the direction of the *Pedobacter* partner, is consistent with a diffusible signal. The signal likely reaches the colony face closest to the *Pedobacter* partner, before appearing to propagate throughout the colony.

Social spreading is dependent upon the presence of a partner. However, the partners in certain combinations of *P. fluorescens* and *Pedobacter* appear to move independently, as we cannot consistently co-isolate both bacteria from all parts of a motile colony in experiments with isolates outside the V48 clade, and even see Pf0-1 move alone when plated distantly from other *Pedobacter* isolates. This first indicates that, in interactions of Pf0-1 with the MAS clade, co-isolation of both participants from the edge of the motile colony may be coincidental, rather than the result of collective movement. This lack of consistency in co-migration also indicates that Pf0-1 is capable of individual “spreading” motility under the conditions explored in this paper, although this movement is still dependent on stimulation by a diffusible signal from the partner species. We propose two possible explanations: either the motility mechanism is distinct from the contact-dependent method of motility seen in interspecies social spreading with *Pedobacter* sp. V48 (14), or the signaling molecule in the V48 clade is tethered to the individual cells, or so closely associated that it necessitates co-migration within the V48 clade to continue induction of movement. In contrast, when close contact is not required to trigger motility, i.e. mediated by a diffusible signal, movement is uncoupled and results in independent migration.

Social assays under high nutrient conditions preclude the social spreading phenotype with strains that are positive for movement on our standard social spreading medium (TSBK-NK), which matches the observations in McCully et al. 2019. Similarly, *Streptomyces* ‘exploratory’ motility triggered by the presence of *Saccharomyces cerevisiae*, can only be induced when a key nutrient is at a low concentration, which occurs in this case once the yeast have depleted the medium of glucose (11). A similar phenomenon has also been observed in *Flavobacterium* individual gliding motility, where decreasing nutrient concentration increases colony diameter, implicating nutrient level as an important factor in regulating gliding motility, and suggesting that nutrient-limiting conditions may activate this movement to colonize surfaces (41–43).

Under low nutrient conditions (10% TSB) the addition of NaCl is required for the interaction between Pf0-1 and V48 (14), but the requirement for this NaCl supplementation is not universal for *P. fluorescens*-*Pedobacter* social spreading. We found that in some interactions lower NaCl favors social spreading, while additional NaCl hinders it, while in other cases the level of NaCl has little influence on the spreading phenotype (Table 2). If the interactions we have explored in this work are environmentally-relevant, the influence of nutritional factors are likely dictated by the predominant conditions of the environment they were isolated from, which could explain the variability in response to salt concentration. Our observations demonstrate that the environment influences interactions between different species, and that there are different responses to variant nutrient conditions, indicating the importance of the role that the environment has in governing the behavioral traits of the system. This finding on the influence of environment on behavior of microbial communities should be considered carefully when deploying small consortia in applications such as plant growth promotion, since the prevailing conditions in different application locations could influence whether desired traits of the introduced consortium are expressed.

As we had hypothesized in our previous report of motility between two distantly-related isolates (14), Interspecies Social Spreading between *P. fluorescens* Pf0-1 and *Pedobacter* sp. V48 does not describe a unique interaction-induced behavior. Rather, it is but an example of a more widespread trait emerging from interactions between members of the *P. fluorescens* group and the *Pedobacter* genus. The fact that many of these bacteria were isolated from distant and diverse environments, suggests that these motility behaviors may be important responses, and could be relevant in an environmental context. The surface spreading phenotypes resulting from these interactions are examples of emergent traits, where the microbial communities have properties that are not found in the individuals. Microbiology has historically focused on the study of individual species, but going forward, it is critical that we consider the context of the entire microbial community, as important activities may result from interactions only be identified when studying bacteria as groups. Although our work uses a simplified two-species model, this is an important step towards learning more about the roles of communities, ultimately working to better understand community behavior as a whole and its interaction with the environment.

## Materials and Methods

### Bacterial strains and culture conditions

Bacterial strains are described in Table 3. Bacteria were routinely grown at 30°C (*Pseudomonas* isolates) or 20°C (*Pedobacter* isolates), in 10% strength tryptic soy broth (BD Difco) amended with NaCl and KH_2_PO_4_, as described by de Boer et al. (44), referred to as TSB-NK (14). To isolate the two species from mixed cultures, we used *Pseudomonas* minimal medium (PMM) with 25 mM succinate (45) for *P. fluorescens* and 14.6 mM lactose for *Pedobacter* as sole carbon sources. Media were solidified with BD Difco Bacto agar (1.5%, wt/vol) when required, except for social spreading assays, for which 2% agar was used. For experiments with variations in nutrients, we used full-strength TSB (30 g/liter), 10% TSB (3 g/liter), and 10% TSB amended with NaCl (TSB-N).

**Table 3.** Bacterial strains.

| **Strain** | **Genotype or Description** | **Source or Reference** |
| --- | --- | --- |
| ***P. fluorescens*** |  |  |
| Pf0-1 | Wild-type isolate from sandy-loam soil, Ap<sup>r</sup><br>CP000094.2 | (34) |
| SBW25 | Isolate from phyllosphere of sugar beet plant NC_012660.1 | (56, 57) |
| AU11518 | Isolate from cystic fibrosis patients<br>NZ_JRXW00000000.1 | (58) |
| AU2989 | Isolate from cystic fibrosis patients<br>NZ_JRXT00000000.1 | (58) |
| AU10973 | Isolate from cystic fibrosis patients<br>Z_JRXV00000000.1 | (58) |
| AU14705 | Isolate from cystic fibrosis patients<br>NZ_JRYA00000000.1 | (58) |
| AU6026 | Isolate from cystic fibrosis patients<br>NZ_JRXU00000000.1 | (58) |
| AU14440 | Isolate from cystic fibrosis patients<br>NZ_JRXX00000000.1 | (58) |
| AU14917 | Isolate from cystic fibrosis patients<br>NZ_JRXY00000000.1 |  |
| A506 | Isolate from pear tree leaves<br>CP003041.1 | (59, 60) |
| Q2-87 | Isolate from wheat rhizosphere<br>CM001558.1 | (60, 61) |
| Q8r1-96 | Isolate from wheat rhizosphere<br>CM001512.1 | (60, 62) |
| F113 | Isolate from root hairs of a sugar-beet plant<br>CP003150.1 | (63, 64) |
| AD21 | Isolate from Dutch coastal dune soil | (33) |
| WH6 | Isolate from wheat rhizosphere (Oregon)<br>NZ_CM001025.1 | Kristen Trippe<br>(65, 66) |
| SF4c | Isolate from wheat rhizosphere (Argentina)<br>JTGH00000000.1 | (67, 68) |
| SF39a | Isolate from wheat rhizosphere (Argentina)<br>JTGG00000000.1 | (67, 68) |
| FW300-N2C3 | Isolate from groundwater (Tennessee)<br>CP012831.1 | (69) |
| FW300-N2E3 | Isolate from groundwater (Tennessee)<br>CP012830.1 | (69) |
| ***P. protegens*** |  |  |
| Pf-5 | Isolate from cotton seedlings<br>NC_004129.6 | (70, 71) |
| ***P. aeruginosa*** |  |  |
| PAO1 | PAO1-L | Dieter Hass;<br>(72) |
| ***Pedobacter*** |  |  |
| V48 | Wild-type isolate from Dutch coastal dune soil<br>AWRU01000031 | (33) |
| *heparinus* DSM<br>2366 | Isolate from dry soil<br>CP001681 | (73, 74) |
| *africanus* DSM<br>12126 | Isolate from soil<br>AJ438171 | (73) |
| *nyackensis* DSM<br>19625 | Isolate from flood plain sediment<br>EU030686 | (75) |
| *agri* PB92 DSM<br>19486 | Isolate from soil<br>EF660751 | (28) |
| *glucosidilyticus* DSM<br>23534 | Isolate from dry riverbed soil<br>EU585748 | (76) |
| *oryzae* N7 DSM<br>19973 | Isolate from rice paddy soil<br>EU109726 | (77) |
| *terricola* DS-45<br>KCTC 12876 | Isolate from soil<br>EF446147 | (32) |
| *panaciterrae* KCTC<br>12594 | Isolate from soil of a ginseng field<br>AB245368 | (78) |
| AL1408 | Isolate from moss rhizosphere | (79) |
| MAS011 | Isolate from sub-Arctic terricolous lichen | (36) |
| MAS018 | Isolate from sub-Arctic terricolous lichen | (36) |
| MAS029 | Isolate from sub-Arctic terricolous lichen | (36) |
| MAS035 | Isolate from sub-Arctic terricolous lichen | (36) |
| G082 | Isolate from sub-Arctic river water | (35) |
| V10001 | Isolate from sub-Arctic river water | (35) |
| JF1508 | Isolate from glacial river water | (80) |
| JF0115 | Isolate from glacial river water | (80) |
| *Pseudopedobacter saltans* DSM12145 | Isolate from soil<br>AJ438173 | (73, 81) |

### Interspecies social spreading assays

Assays were performed as described in McCully et al (2019). Briefly, *Pseudomonas* and *Pedobacter* isolates were grown in liquid TSB-NK at 20°C overnight. Assays were performed on TSB-NK solidified with 2% agar.

### (i) Mixed inoculum assays

Five microliters of each participant were combined in one spot on the agar surface. After inoculation liquid had dried, plates were incubated at 20°C. Measurements of the colony diameter were taken every 24 h. Experiments were performed in triplicate.

### (ii) Diffusion/signaling assay—adjacent plating

*P. fluorescens* and *Pedobacter* isolates were grown as described above. The aliquots of bacteria were plated adjacent but without the drops touching at distances of 5 mm, 7.5 mm, 10 mm, 11 mm, 12 mm, 13 mm, 14 mm, 15 mm and 20 mm from the centers of each colony. Once the inoculation liquid had dried, plates were incubated at 20°C and monitored daily to determine the time at which colony growth led to contact between the isolates, and/or when spreading phenotypes developed.

### (iii) Cellophane overlay assay

Squares of porous cellophane (GE Healthcare Biosciences Corp.) were placed on top of TSB-NK plates. Cultures of various *Pedobacter* isolates were placed on top of the cellophane to allow metabolites to diffuse into the agar, with cellophane alone used as a negative control. Plates were incubated at 20°C for 2 days, at which point cellophane was removed, and 5 µl spots of *P. fluorescens* Pf0-1 were placed in the center of the plate. Pf0-1 was placed on plates where cellophane had been (negative control), or plates where the partner species had been cultured on top of the cellophane. Measurements of the colony diameter were taken every 24 h. Experiments were performed in triplicate.

### (iv) Volatile signaling

Ten microliter aliquots of cultures of *Pedobacter* MAS isolates and Pf0-1 were plated onto separate TSB-NK plates. A Petri dish ‘sandwich’ was made by pairing the bottom of each inoculated Petri dish (one with Pf0-1 and one with a *Pedobacter* isolate), and held together with parafilm. Measurements of the colony diameter were taken every 24 h. Experiments were performed in triplicate.

### 16S rRNA gene sequencing

*Pedobacter* colonies were picked using sterile toothpicks, suspended in 25 µl colony lysis buffer (1% Triton X-100 in 0.02 mol l-1 Tris / 0.002 mol l-1 EDTA at pH 8.0), heated to 95°C for 10 min, and then cooled to 4 °C. From the resulting lysate, 1 µL was used as a template in a 25μL PCR reaction (35 cycles, annealing at 51 °C for 30 s, extension at 68 °C for 90 s, denaturing at 95 °C for 30 s) in an MJR PTC-200 thermocycler (MJ Research Inc.) using primers 8F (5’-AGAGTTTGATCCTGGCTCAG-3’) and 1522R (5’-AAGGAGGTGATCCAACCGCA-3’) and 0.75 units of Taq polymerase (New England Biolabs) (46, 47). Amplicons were visualized on an 0.8% agarose gel using SYBR Safe (Life Technologies) and purified for DNA sequencing using 20 units of exonuclease I and 5 units of Antarctic phosphatase (New England Biolabs) at 37 °C for 30 min. Enzymes were deactivated at 90 °C for 5 min. Partial sequencing of purified amplicons were primed using primers 519F (5’-CAGCAGCCGCGGTAATAC-3’) and 926R (5’-CCGTCAATTCCTTTGAGTTT-3’), or 8F (46, 48). The BigDye terminator kit was used to generate products for sequencing with an Applied Biosystems 3130XL DNA analyzer (Applied Biosystems) at Macrogen Europe, Amsterdam, the Netherlands. Sequences were examined in ABI Sequence Scanner 1.0 (Applied Biosystems) and the forward sequence and the reverse complement of the reverse sequence were manually aligned and merged.

### Phylogenetic analysis

The 16S rRNA gene sequence was used to evaluate relatedness between *Pedobacter* isolates. Sequences were obtained from the SILVA (https://www.arb-silva.de/) database (49), or determined as described above. To evaluate relatedness of *Pseudomonas* sp., the sequences of housekeeping genes *gapA, gltA, gyrB, and rpoB* were obtained from complete or draft genome sequences of each isolate used. Alignments for each gene were created using Muscle (50) after which sequences were trimmed to equal lengths. The trimmed housekeeping genes for each strain were then concatenated. Genome sequences were accessed using pseudomonas.com (51).

The evolutionary histories for *Pedobacter* and *Pseudomonas* sp. were inferred by using the Maximum Likelihood method based on the General Time Reversible model (17) using MEGA7 (18). Sequences for phylogenetic analysis of each genus were aligned using ClustalW (52). To determine the most appropriate phylogenetic model, the FindModel tool was used (https://www.hiv.lanl.gov/content/sequence/findmodel/findmodel.html) (53). For the molecular phylogenetic analysis, bootstrap consensus trees were inferred from 1000 replicates and are taken to represent the evolutionary history of the taxa analyzed (54). Branches corresponding to partitions reproduced in less than 50% bootstrap replicates are collapsed. Initial tree(s) for the heuristic search were obtained automatically by applying Neighbor-Join and BioNJ algorithms to a matrix of pairwise distances estimated using the Maximum Composite Likelihood (MCL) approach, and then selecting the topology with superior log likelihood value. A discrete Gamma distribution was used to model evolutionary rate differences among sites (4 categories (+G, parameter = 0.2139 for *Pedobacter* sp. and 0.2333 for *Pseudomonas* sp.)). For the *Pedobacter* sp. analysis there were a total of 1547 positions in the final dataset, while for *Pseudomonas* sp. there were 8635 positions in the final dataset. Phylogenetic trees were annotated in iTOL (https://itol.embl.de/) (55).

### Imaging

Still pictures were taken using an EOS Rebel T3i camera (Canon) and a Google Pixel 2 and processed using Photoshop CC 2017 version: 14.2.1 and GIMP (GNU Image Manipulation Program) version: 2.10.10. Using Photoshop, the levels of some images were adjusted to improve contrast.

### Statistics

For cellophane overlay assays, we measured the amount of colony expansion of *P. fluorescens* Pf0-1 to test induction of social spreading by diffusible signals, after preconditioning the agar by growing monocultures of *Pedobacter* isolates, or with sterile cellophane as a control. For volatile signaling assays, we measured the amount of colony expansion of the monocultures of *P. fluorescens* Pf0-1 exposed to *Pedobacter* isolates, as well as an unexposed control, in a Petri dish sandwich. In each assay, colony diameter of three independent experiments was measured every 24 h. Changes in colony diameter from the control were evaluated by using an ANOVA with a Tukey HSD post-hoc test, in R Studio 3.5.2.

## References

1. Philippot L, Raaijmakers JM, Lemanceau P, van der Putten WH. 2013. Going back to the roots: The microbial ecology of the rhizosphere. Nat Rev Microbiol 11:789–799.

2. Fierer N. 2017. Embracing the unknown: disentangling the complexities of the soil microbiome. Nat Rev Microbiol 15:579–590.

3. Van Der Heijden MGA, Bardgett RD, Van Straalen NM. 2008. The unseen majority: Soil microbes as drivers of plant diversity and productivity in terrestrial ecosystems. Ecol Lett 11:296–310.

4. Wall DH, Nielsen UN, Six J. 2015. Soil biodiversity and human health. Nature 528:69–76.

5. Rajkumar M, Sandhya S, Prasad MNV, Freitas H. 2012. Perspectives of plant-associated microbes in heavy metal phytoremediation. Biotechnol Adv 30:1562–1574.

6. Hall EK, Bernhardt ES, Bier RL, Bradford MA, Boot CM, Cotner JB, del Giorgio PA, Evans SE, Graham EB, Jones SE, Lennon JT, Locey KJ, Nemergut D, Osborne BB, Rocca JD, Schimel JP, Waldrop MP, Wallenstein MD. 2018. Understanding how microbiomes influence the systems they inhabit. Nat Microbiol 3:977–982.

7. Konopka A. 2009. What is microbial community ecology? ISME J 3:1223–1230.

8. Madsen JS, Sørensen SJ, Burmølle M. 2018. Bacterial social interactions and the emergence of community-intrinsic properties. Curr Opin Microbiol 42:104–109.

9. Stubbendieck RM, Straight PD. 2015. Escape from lethal bacterial competition through coupled activation of antibiotic resistance and a mobilized subpopulation. PLoS Genet 11:e1005722.

10. Liu Y, Kyle S, Straight PD. 2018. Antibiotic stimulation of a *Bacillus subtilis* migratory response. mSphere 3:e00586–17.

11. Jones SE, Ho L, Rees CA, Hill JE, Nodwell JR, Elliot MA. 2017. Streptomyces exploration is triggered by fungal interactions and volatile signals. eLife 6:e21738.

12. Limoli DH, Donegan NP, Warren EA, Cheung AL, O’toole GA. 2019. Interspecies signaling generates exploratory motility in Pseudomonas aeruginosa. bioRxiv.

13. Hagai E, Dvora R, Havkin-Blank T, Zelinger E, Porat Z, Schulz S, Helman Y. 2014. Surface-motility induction, attraction and hitchhiking between bacterial species promote dispersal on solid surfaces. ISME J 8:1147–1151.

14. McCully LM, Bitzer AS, Seaton SC, Smith LM, Silby MW. 2019. Interspecies Social Spreading: Interaction between two sessile soil bacteria leads to emergence of surface motility. mSphere 4:1–16.

15. Venturi V, Bertani I, Kerényi Á, Netotea S, Pongor S. 2010. Co-swarming and local collapse: Quorum sensing conveys resilience to bacterial communities by localizing cheater mutants in *Pseudomonas aeruginosa*. PLoS One 5:e9998.

16. Samad T, Billings N, Birjiniuk A, Crouzier T, Doyle PS, Ribbeck K. 2017. Swimming bacteria promote dispersal of non-motile staphylococcal species. ISME J 11:1933–1937.

17. Nei M, Kumar S. 2000. Molecular Evolution and Phylogenetics 1 edition. Oxford University Press.

18. Kumar S, Stecher G, Tamura K. 2016. MEGA7: Molecular Evolutionary Genetics Analysis Version 7.0 for Bigger Datasets. Mol Biol Evol 33:1870–1874.

19. Garbeva P, Silby MW, Raaijmakers JM, Levy SB, de Boer W. 2011. Transcriptional and antagonistic responses of *Pseudomonas fluorescens* Pf0-1 to phylogenetically different bacterial competitors. ISME J 5:973–985.

20. Garbeva P, Tyc O, Remus-Emsermann MNP, van der Wal A, Vos M, Silby M, de Boer W. 2011. No apparent costs for facultative antibiotic production by the soil bacterium *Pseudomonas fluorescens* Pf0-1. PLoS One 6:e27266.

21. Garbeva P, Hordijk C, Gerards S, de Boer W. 2014. Volatile-mediated interactions between phylogenetically different soil bacteria. Front Microbiol 5:1–9.

22. Finkelshtein A, Roth D, Ben Jacob E, Ingham CJ. 2015. Bacterial swarms recruit cargo bacteria to pave the way in toxic environments. mBio 6:e00074–15.

23. Ingham CJ, Kalisman O, Finkelshtein A, Ben-Jacob E. 2011. Mutually facilitated dispersal between the nonmotile fungus *Aspergillus fumigatus* and the swarming bacterium *Paenibacillus vortex*. Proc Natl Acad Sci 108:19731–19736.

24. Poehlein A, Daniel R, Simeonova DD. 2015. Genome sequence of *Pedobacter glucosidilyticus* DD6b, isolated from zooplankton Daphnia magna. Stand Genomic Sci 10:10.1186/s40793-015-0086-x.

25. Kang H, Kim H, Joung Y, Joh K. 2014. Pedobacter rivuli sp. nov., isolated from a freshwater stream. Int J Syst Evol Microbiol 64:4073–4078.

26. Oh HW, Kim BC, Park DS, Jeong WJ, Kim H, Lee KH, Kim SU. 2013. Pedobacter luteus sp. nov., isolated from soil. Int J Syst Evol Microbiol 63:1304–1310.

27. Kwon SW, Son JA, Kim SJ, Kim YS, Park IC, Bok JI, Weon HY. 2011. *Pedobacter rhizosphaerae* sp. nov. and *Pedobacter soli* sp. nov., isolated from rhizosphere soil of Chinese cabbage (*Brassica campestris*). Int J Syst Evol Microbiol 61:2874–2879.

28. Roh SW, Quan ZX, Nam Y Do, Chang HW, Kim KH, Kim MK, Im WT, Jin L, Kim SH, Lee ST, Bae JW. 2008. *Pedobacter agri* sp. nov., from soil. Int J Syst Evol Microbiol 58:1640–1643.

29. Švec P, Králová S, Busse HJ, Kleinhagauer T, Kýrová K, Pantůček R, Mašlaňová I, Staňková E, Némec M, Holochová P, Barták M, Sedláček I. 2017. *Pedobacter psychrophilus* sp. nov., isolated from fragmentary rock. Int J Syst Evol Microbiol 67:2538–2543.

30. Wang Z, Tan Y, Xu D, Wang G, Yuan J, Zheng S. 2016. Pedobacter vanadiisoli sp. nov., isolated from soil of a vanadium mine. Int J Syst Evol Microbiol 66:5112–5117.

31. Yuan K, Cao M, Li J, Wang G. 2018. *Pedobacter mongoliensis* sp. nov., isolated from grassland soil. Int J Syst Evol Microbiol 68:1112–1117.

32. Yoon J-H, Kang S-J, Park S, Oh T-K. 2007. *Pedobacter lentus* sp. nov. and *Pedobacter terricola* sp. nov., isolated from soil. Int J Syst Evol Microbiol 57:2089–2095.

33. de Boer W, Verheggen P, Klein Gunnewiek PJA, Kowalchuk GA, van Veen JA. 2003. Microbial community composition affects soil fungistasis. Appl Environ Microbiol 69:835–844.

34. Compeau G, Al-Achi BJ, Platsouka E, Levy SB. 1988. Survival of rifampin-resistant mutants of *Pseudomonas fluorescens* and *Pseudomonas putida* in soil systems. Appl Environ Microbiol 54:2432–2438.

35. Markúsdóttir M, Heidmarsson S, Eythórsdóttir A, Magnússon KP, Vilhelmsson O. 2013. The natural and anthropogenic microbiota of Glerá, a sub-arctic river in northeastern Iceland. Int Biodeterior Biodegrad 84:192–203.

36. Sigurbjörnsdóttir MA, Vilhelmsson O. 2016. Selective isolation of potentially phosphate-mobilizing, biosurfactant-producing and biodegradative bacteria associated with a sub-Arctic, terricolous lichen, *Peltigera membranacea*. FEMS Microbiol Ecol 92:10.1093/femsec/fiw090.

37. Brabcová V, Nováková M, Davidová A, Baldrian P. 2016. Dead fungal mycelium in forest soil represents a decomposition hotspot and a habitat for a specific microbial community. New Phytol 210:1369–1381.

38. Männistö MK, Häggblom MM. 2006. Characterization of psychrotolerant heterotrophic bacteria from Finnish Lapland. Syst Appl Microbiol 29:229–243.

39. Thakur V, Kumar V, Kumar S, Singh D. 2018. Diverse culturable bacterial communities with cellulolytic potential revealed from pristine habitat in Indian trans-Himalaya. Can J Microbiol 64:798–808.

40. Zhu X, Jin L, Sun K, Li S, Li X, Ling W. 2016. Phenanthrene and pyrene modify the composition and structure of the cultivable endophytic bacterial community in ryegrass (*Lolium multiflorum* Lam). Int J Environ Res Public Health 13.

41. Laanto E, Bamford JKH, Laakso J, Sundberg LR. 2012. Phage-driven loss of virulence in a fish pathogenic bacterium. PLoS One 7:e53157.

42. Pérez-Pascual D, Menéndez A, Fernández L, Méndez J, Reimundo P, Navais R, Guijarro JA. 2009. Spreading versus biomass production by colonies of the fish pathogen *Flavobacterium psychrophilum*: Role of the nutrient concentration. Int Microbiol 12:207–214.

43. Penttinen R, Hoikkala V, Sundberg LR. 2018. Gliding motility and expression of motility-related genes in spreading and non-spreading colonies of *Flavobacterium columnare*. Front Microbiol 9:1–12.

44. De Boer W, Wagenaar AM, Klein Gunnewiek PJA, Van Veen JA. 2007. In vitro suppression of fungi caused by combinations of apparently non-antagonistic soil bacteria. FEMS Microbiol Ecol 59:177–185.

45. Kirner S, Krauss S, Sury G, Lam ST, Ligon JM, van Pee K-H. 1996. The non-haem chloroperoxidase from *Pseudomonas fluorescens* and its relationship to pyrrolnitrin biosynthesis. Microbiology 142:2129–2135.

46. Turner S, Pryer KM, Miao VP, Palmer JD. 1999. Investigating deep phylogenetic relationships among cyanobacteria and plastids by small subunit rRNA sequence analysis. J Eukaryot Microbiol 46:327–338.

47. Giovannoni SJ. 1991. The polymerase chain reaction., p. 177–201. *In* Stackebrandt, E, Goodfellow, M (eds.), Sequencing and Hybridization Techniques in Bacterial Systematics. Wiley, New York.

48. Lane DJ. 1991. 16S/23S rRNA sequencing, p. 115–175. *In* Stackebrandt, E, Goodfellow, M (eds.), Nucleic acid techniques in bacterial systematics. John Wiley and Sons, New York, NY.

49. Quast C, Pruesse E, Yilmaz P, Gerken J, Schweer T, Yarza P, Peplies J, Glöckner FO. 2013. The SILVA ribosomal RNA gene database project: Improved data processing and web-based tools. Nucleic Acids Res 41:590–596.

50. Edgar RC. 2004. MUSCLE: multiple sequence alignment with high accuracy and high throughput. Nucleic Acids Res 32:1792–7.

51. Winsor GL, Griffiths EJ, Lo R, Dhillon BK, Shay JA, Brinkman FSL. 2016. Enhanced annotations and features for comparing thousands of *Pseudomonas* genomes in the Pseudomonas genome database. Nucleic Acids Res 44:D646–D653.

52. Larkin MA, Blackshields G, Brown NP, Chenna R, Mcgettigan PA, McWilliam H, Valentin F, Wallace IM, Wilm A, Lopez R, Thompson JD, Gibson TJ, Higgins DG. 2007. Clustal W and Clustal X version 2.0. Bioinformatics 23:2947–2948.

53. Posada D, Crandall KA. 1998. MODELTEST: Testing the model of DNA substitution. Bioinformatics 14:817–818.

54. Felsenstein J. 1985. Confidence limits on phylogenies: An approach using the bootstrap. Evolution (N Y) 39:783–791.

55. Letunic I, Bork P. 2016. Interactive tree of life (iTOL) v3: an online tool for the display and annotation of phylogenetic and other trees. Nucleic Acids Res 44:W242–W245.

56. Rainey PB, Bailey MJ. 1996. Physical and genetic map of the *Pseudomonas fluorescens* SBW25 chromosome. Mol Microbiol 19:521–533.

57. Thompson IP, Lilley AK, Ellis RJ, Bramwell PA, Bailey MJ. 1995. Survival, colonization and dispersal of genetically modified *Pseudomonas fluorescens* SBW25 in the phytosphere of field grown sugar beet. Nat Biotechnol 13:1493–1497.

58. Scales BS, Erb-Downward JR, LiPuma JJ, Huffnagle GB. 2015. Draft genome sequences of five *Pseudomonas fluorescens* subclade I and II strains, isolated from human respiratory samples. Genome Announc 3:2012–2013.

59. Wilson M, Lindow SE. 1993. Interactions between the biological control agent *Pseudomonas fluorescens* A506 and *Erwinia amylovora* in pear blossoms. Ecol Epidemiol 83:117–123.

60. Loper JE, Hassan KA, Mavrodi D V., Davis EW, Lim CK, Shaffer BT, Elbourne LDH, Stockwell VO, Hartney SL, Breakwell K, Henkels MD, Tetu SG, Rangel LI, Kidarsa TA, Wilson NL, van de Mortel JE, Song C, Blumhagen R, Radune D, Hostetler JB, Brinkac LM, Durkin AS, Kluepfel DA, Wechter WP, Anderson AJ, Kim YC, Pierson LS, Pierson EA, Lindow SE, Kobayashi DY, Raaijmakers JM, Weller DM, Thomashow LS, Allen AE, Paulsen IT. 2012. Comparative genomics of plant-associated *Pseudomonas* spp.: Insights into diversity and inheritance of traits involved in multitrophic interactions. PLoS Genet 8.

61. Vincent MN, Harrison LA, Brackin JM, Kovacevich PA, Mukerji P, Weller DM, Pierson EA. 1991. Genetic analysis of the antifungal activity of a soilborne *Pseudomonas aureofaciens* strain. Appl Environ Microbiol 57:2928–2934.

62. Raaijmakers JM, Weller DM. 1998. Natural plant protection by 2,4-diacetylphloroglucinol-producing *Pseudomonas* spp. in Take-all decline soils. Mol Plant-Microbe Interact 11:144–152.

63. Redondo-Nieto M, Matthieu B, Morrisey JP, Germaine K, Martínez Granero F, Barahona E, Navazo A, Sánchez Contreras M, Moynihan JA, Giddens SR, Coppoolse ER, Candela M, Stiekema WJ, Rainey PB, Dowling D, O’Gara F, Martín M, Rivilla R. 2012. Genome sequence of the biocontrol strain *Pseudomonas fluorescens* F113. J Bacteriol 194:1273–1274.

64. Shanahan P, O’Sullivan DJ, Simpson P, Glennon JD, O’Gara F. 1992. Isolation of a 2,4- diacetylphloroglucinol from a fluorescent pseudomonad and investigation of physiological parameters influencing its production. Appl Environ Microbiol 58:353–358.

65. Kimbrel JA, Givan SA, Halgren AB, Creason AL, Mills DI, Banowetz GM, Armstrong DJ, Chang JH. 2010. An improved, high-quality draft genome sequence of the Germination-Arrest Factor-producing *Pseudomonas fluorescens* WH6. BMC Genomics 11:522.

66. Banowetz GM, Azevedo MD, Armstrong DJ, Halgren AB, Mills DI. 2008. Germination-Arrest Factor (GAF): Biological properties of a novel, naturally-occurring herbicide produced by selected isolates of rhizosphere bacteria. Biol Control 46:380–390.

67. Fischer SE, Fischer SI, Magris S, Mori GB. 2007. Isolation and characterization of bacteria from the rhizosphere of wheat. World J Microbiol Biotechnol 23:895–903.

68. Ly LK, Underwood GE, McCully LM, Bitzer AS, Godino A, Bucci V, Brigham CJ, Príncipe A, Fischer SE, Silby MW. 2016. Draft genome sequences of *Pseudomonas fluorescens* strains SF39a and SF4c, potential plant growth promotion and biocontrol agents. Genome Announc 3:e00219–15.

69. Price MN, Wetmore KM, Waters RJ, Callaghan M, Ray J, Liu H, Kuehl J V, Melnyk RA, Lamson JS, Suh Y, Carlson HK, Esquivel Z, Sadeeshkumar H, Chakraborty R, Zane GM, Rubin BE, Wall JD, Visel A, Bristow J, Blow MJ, Arkin AP, Deutschbauer AM. 2018. Mutant phenotypes for thousands of bacterial genes of unknown function. Nature 557:503–509.

70. Howell CR, Stipanovic RD. 1979. Control of *Rhizoctonia solani* on cotton seedlings with *Pseudomonas fluorescens* and with an antibiotic produced by the bacterium. Phytopathology 69:480–482.

71. Paulsen IT, Press CM, Ravel J, Kobayashi DY, Myers GSA, Mavrodi D V, DeBoy RT, Seshadri R, Ren Q, Madupu R, Dodson RJ, Durkin AS, Brinkac LM, Daugherty SC, Sullivan SA, Rosovitz MJ, Gwinn ML, Zhou L, Schneider DJ, Cartinhour SW, Nelson WC, Weidman J, Watkins K, Tran K, Khouri H, Pierson EA, Pierson LS, Thomashow LS, Loper JE. 2005. Complete genome sequence of the plant commensal *Pseudomonas fluorescens* Pf-5. Nat Biotechnol 23:873–878.

72. Heurlier K, Dénervaud V, Pessi G, Reimmann C, Haas D. 2003. Negative control of quorum sensing by RpoN (s54) in *Pseudomonas aeruginosa* PAO1. J Bacteriol 185:2227–2235.

73. Steyn PL, Segers P, Vancanneyt M, Sandra P, Kersters K, Joubert JJ. 1998. Classification of heparinolytic bacteria into a new genus, *Pedobacter*, comprising four species: *Pedobacter heparinus* comb. nov., *Pedobacter piscium* comb. nov., *Pedobacter africanus* sp. nov. and *Pedobacter saltans* sp. nov. Int J Syst Bacteriol 48:165–177.

74. Payza AN, Korn ED. 1956. The degradation of heparin by bacterial enzymes I. Adaptation and lyophilized cells. J Biol Chem 223:853–858.

75. Gordon NS, Valenzuela A, Adams SM, Ramsey PW, Pollock JL, Holben WE, Gannon JE. 2009. *Pedobacter nyackensis* sp. nov., *Pedobacter alluvionis* sp. nov. and *Pedobacter borealis* sp. nov., isolated from Montana flood-plain sediment and forest soil. Int J Syst Evol Microbiol 59:1720–1726.

76. Luo X, Wang Z, Dai J, Zhang L, Li J, Tang Y, Wang Y, Fang C. 2010. Pedobacter glucosidilyticus sp. nov., isolated from dry riverbed soil. Int J Syst Evol Microbiol 60:229–233.

77. Jeon Y, Kim JM, Park JH, Lee SH, Seong CN, Lee SS, Jeon CO. 2009. Pedobacter oryzae sp. nov., isolated from rice paddy soil. Int J Syst Evol Microbiol 59:2491–2495.

78. Yoon M-H, Ten LN, Im W-T, Lee S-T. 2007. Pedobacter panaciterrae sp. nov., isolated from soil in South Korea. Int J Syst Evol Microbiol 57:381–386.

79. Brynjarsdóttir ÁI. 2013. Söfnun, einangrun og tegundagreining bakteríustofna úr íslenskum jarðvegsgerðum [Sampling, isolation and identification of bacterial strains from Icelandic soil types]. University of Akureyri.

80. Jóelsson JP, Frijónsdóttir H, Vilhelmsson O. 2013. Bioprospecting a glacial river in Iceland for bacterial biopolymer degraders. Cold Reg Sci Technol 96:86–95.

81. Cao J, Lai Q, Li G, Shao Z. 2014. *Pseudopedobacter beijingensis* gen. nov., sp. nov., isolated from coking wastewater activated sludge, and reclassification of *Pedobacter saltans* as *Pseudopedobacter saltans* comb. nov. Int J Syst Evol Microbiol 64:1853–1858.

